# Parallel evolution of amphioxus and vertebrate small-scale gene duplications

**DOI:** 10.1101/2022.01.18.476203

**Authors:** Marina Brasó-Vives, Ferdinand Marlétaz, Amina Echchiki, Federica Mantica, Rafael D. Acemel, José L. Gómez-Skarmeta, Diego A. Hartasánchez, Lorlane L. Targa, Pierre Pontarotti, Juan J. Tena, Ignacio Maeso, Hector Escriva, Manuel Irimia, Marc Robinson-Rechavi

## Abstract

**Background:** Amphioxus are non-vertebrate chordates characterized by a slow morphological and molecular evolution. They share the basic chordate body-plan and genome organization with vertebrates but lack their 2*R* whole-genome duplications and their developmental complexity. For these reasons, amphioxus are frequently used as an outgroup to study vertebrate genome evolution and Evo-Devo. Aside from whole-genome duplications, genes continuously duplicate on a smaller scale. Small-scale duplicated genes can be found in both amphioxus and vertebrate genomes, while only the vertebrate genomes have duplicated genes product of their 2R whole-genome duplications. Here, we explore the history of small-scale gene duplications in the amphioxus lineage and compare it to small- and large-scale gene duplication history in vertebrates.

**Results:** We present a study of the European amphioxus (*Branchiostoma lanceolatum*) gene duplications thanks to a new, high-quality genome reference. We find that, despite its overall slow molecular evolution, the amphioxus lineage has had a history of small-scale duplications similar to the one observed in vertebrates. We find parallel gene duplication profiles between amphioxus and vertebrates, and conserved functional constraints in gene duplication. Moreover, amphioxus gene duplicates show levels of expression and patterns of functional specialization similar to the ones observed in vertebrate duplicated genes. We also find strong conservation of gene synteny between two distant amphioxus species, *B. lanceolatum* and *B. floridae*, with two major chromosomal rearrangements.

**Conclusions:** In contrast to their slower molecular and morphological evolution, amphioxus’ small-scale gene duplication history resembles that of the vertebrate lineage both in quantitative and in functional terms.

## Background

Amphioxus, extant members of the subphylum Cephalochordata, are small marine non-vertebrate chordates key to the study of chordate and early vertebrate evolution [1–3]. Cephalochordates, together with tunicates, are the only two known extant non-vertebrate chordate lineages. In contrast to tunicates, which present very derived genomes and adult morphology, the amphioxus lineage has had a slower molecular evolution and presents ancestral phenotypic characters both during development and in adult individuals [4–7]. For these reasons, amphioxus have been historically used as outgroup models to study vertebrate anatomy, embryonic development, and genomics [6]. Comparative genomics of amphioxus and other chordates has been key to understanding vertebrate genome and transcriptome origins [6–10] and continues to shed light on the ancestral genome structure of chordates [11].

Among other findings, the study of amphioxus genomes validated the *2R* hypothesis proposing two rounds of whole-genome duplication (WGD) in early vertebrate evolution [6,9,12]. Duplication, both whole-genome and small-scale, is an important contributor to genome evolution [13–18]. Gene duplication has been widely studied within vertebrates, which show a great diversity of duplication rates, including both WGDs (2R and fish- or amphibian-specific) and small-scale duplications [18–22]. Notably, all vertebrates share an abundant set of gene duplicates derived from the two “2R” rounds of WGD, the 2R ohnologs. Additionally to these 2R ohnologs, small-scale duplications have been frequent in the vertebrate lineage since its early evolution, and are well studied. In contrast, little is known about gene duplication in the amphioxus lineage, except for the lack of WGDs [6]. Given the difference in the origin of duplicated genes in both lineages, and the important contribution that 2R ohnologs have in the vertebrate gene pool, the study of gene duplications in both amphioxus and vertebrates is key to understanding the differences in gene duplication history between these lineages, and the role of duplication in both lineages’ evolution.

Studying gene duplication in amphioxus is complicated by their very high heterozygosity. Amphioxus present one of the highest ever measured heterozygosity rates in animals, exceeded by few others such as the purple sea urchin *Strongylocentrotus purpuratus* [6,23,24]. When building a genome assembly, high heterozygosity can result in alternative haplotypes assembled as separate loci [25,26], and in difficulty distinguishing between alleles and recent duplications. Duplicated sequences must neither be collapsed into a single region nor placed as alternative haplotypes [27,28]. Thus, assembling the genome with long-read and long-distance data, as well as using appropriate haplotype collapsing methods, is critical to the study of gene duplications in amphioxus [11].

The European amphioxus (*Branchiostoma lanceolatum*) is one of the best studied species of amphioxus. It has an ecological range extending at least from the northeastern Atlantic Ocean to the Mediterranean Sea [1] and a diploid karyotype of 19 pairs of chromosomes [29]. For this study, we built the first long-read and proximity-ligation based high-quality genome assembly, and we improved gene annotation. Thanks to the reliability provided by a good-quality genome reference and annotation, we present here an analysis of the gene duplication evolution of the lineage leading to the European amphioxus. We compared the amphioxus gene duplication history to that of vertebrates, with special focus on general vertebrate trends, such as the gene duplicates retained after the 2R WGDs (2R ohnologs). This work represents the first detailed study of gene duplication history in the amphioxus lineage compared to the vertebrate lineage, and sheds light on the role of gene duplication in the early evolution of chordates. Given the presence of the 2R WGDs in the vertebrate lineage and the absence of WGD in the amphioxus lineage, and the overall slower evolutionary rate of amphioxus genomes, one would expect to find substantial differences in the history of gene duplications during evolution of these two lineages. Surprisingly, we find conserved gene duplication patterns and parallel functional constraints in gene duplication between the two lineages.

## Results

### High-quality *B. lanceolatum* genome assembly and annotation

We constructed BraLan3, a high-quality genome assembly combining PacBio sequencing reads with chromosome conformation Hi-C data of the European amphioxus, *B. lanceolatum* (see Methods). We re-annotated and validated protein-coding genes in this new genome reference (see Methods). Basic genome assembly quality statistics portray the high-quality of BraLan3 (Figure 1, Table 1). It has an N50 of 23,752,511 bp for a total length of 474,791,770 bp. Moreover, 96.78% of the BraLan3 sequence is found within the 19 chromosomes of *B. lanceolatum’s* haploid karyotype [29]. The gene annotation of BraLan3 contains 27,102 protein coding genes of which 96.97% (26,282 genes) are in chromosomes and 97.66% (26,468 genes) are supported by strong evidence (see Methods). This new annotation has a BUSCO completeness score of 97.6%. These numbers represent a strong improvement in genome assembly and gene annotation quality for *B. lanceolatum* relative to the previous short-read based genome [8]. The quality of BraLan3 resembles that of highly studied vertebrate species, and is as good or better than those of other amphioxus species genomes in all measured statistics (Figure 1, Figure S1, Table 1).

**Figure 1.**
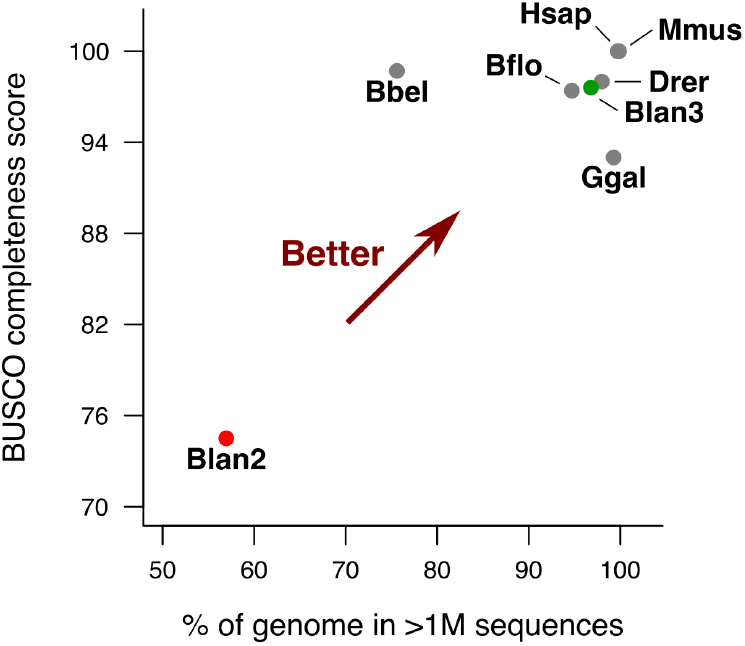
BraLan3 genome assembly and annotation quality comparison. The x axis represents the percentage of a genome assembly sequence in chromosomal-sized sequences (>1M nucleotides). The y axis represents the BUSCO completeness score of a genome annotation. Each data point corresponds to a genome assembly: Drer corresponds to the genome assembly for zebrafish (GRCz11), Ggal for chicken (GRCg6a), Mmus for mouse (GRCm39), Hsap for human (GRCh38), Bflo for an American amphioxus (*B.floridae*) [9], Bbel for an Asian amphioxus (*B. belcheri*) [7]; Blan2 corresponds to the previously available genome reference for the European amphioxus [8]; and Blan3 corresponds to BraLan3, the genome reference for the European amphioxus presented in this study. BUSCO completeness score was performed with the metazoan gene universe in all cases. Table 1 contains this figure’s data and additional genome statistics.

**Table 1.** BraLan3 assembly and annotation quality comparison. All statistics were calculated with the chromosome-level assembly when available or, alternatively, with the scaffold-level assembly; scaffold vs. chromosome-level especially impacts N50 and L50. BUSCO completeness score was performed with the metazoan gene universe in all cases. BraLan3 corresponds to the assembly presented in this work, BraLan2 refers to the previously available *B. lanceolatum* assembly [8].

|  | Total length (Mbp) | N50 (Mbp) | L50 | # sequences >1Mbp | % length in >1Mbp | # gaps in >1Mbp | BUSCO completeness |
| --- | --- | --- | --- | --- | --- | --- | --- |
| **BraLan3** | 474.79 | 23.75 | 8 | 19 | 96.78% | 1523 | 97.60% |
| **BraLan2** | 495.35 | 1.30 | 79 | 108 | 56.95% | 54787 | 74.50% |
| ***B. floridae*** | 513.46 | 25.44 | 9 | 20 | 94.70% | 20731 | 97.40% |
| ***B. belcheri*** | 426.12 | 2.33 | 52 | 121 | 75.60% | 12649 | 98.70% |
| **Zebrafish (GRCz11)** | 1,373.45 | 54.30 | 11 | 25 | 97.96% | 17943 | 98.00% |
| **Chicken (GRCg6a)** | 1,065.35 | 91.32 | 4 | 37 | 99.27% | 866 | 93.00% |
| **Human (GRCh38)** | 3,099.73 | 145.14 | 9 | 25 | 99.71% | 997 | 100.00% |
| **Mouse (GRCm39)** | 2,728.21 | 130.53 | 9 | 21 | 99.86% | 268 | 100.00% |

As an example of the completeness of the assembly, we were able to identify for the first time in this species’ genome assembly three *ProtoRAG* transposon sequences. The recombination-activating genes (RAG1 and RAG2) in jawed vertebrates, essential for V(D)J recombination in immunoglobulin genes, have their origin in a transposon domestication in vertebrate evolution, a RAGB transposon. An active *ProtoRAG* transposon has been described in the *B. belcheri* genome containing both RAG1 and RAG2 genes flanked by terminal inverted repeats (TIR) [30]. We screened BraLan3 for the presence of *ProtoRAG* transposons and found two full *ProtoRAG* transposon copies in chromosomes 12 and 18 (with 98.8% sequence similarity) and an incomplete copy with a truncated RAG1 in chromosome 16 (Supplementary Note 1). These *B. lanceolatum ProtoRAG* transposons present the same structure as the *ProtoRAG* of *B. belcheri,* that is, without a PHD zinc-finger domain or plant homeodomain in RAG2, unlike in jawed vertebrates [31]. In addition, we found 13 Miniature Inverted-repeat Transposable Elements (MITE) in 10 different BraLan3 chromosomes, suggesting that the *ProtoRAG* transposon is active in *B. lanceolatum*. Together, these results show that this transposon has been active in amphioxus at least since the split between the *B. belcheri* lineage and the *B. lanceolatum* lineage.

### Gene duplication profiles in amphioxus and vertebrates

In order to study the gene duplication history of *B. lanceolatum*, we inferred ortholog gene groups (orthogroups) for chordates using the new BraLan3 gene annotation. We define an orthogroup as a set of genes derived from a single gene in the chordate last common ancestor. We included in the orthogroup analysis two other amphioxus species, Florida amphioxus (*B. floridae*) and an Asian amphioxus (*B. belcheri*), and four vertebrate species: zebrafish (*D. rerio*), chicken (*G. gallus*), mouse (*M. musculus*) and human (*H. sapiens*). We did not include ascidian genomes due to their highly derived status [2], which makes them less useful to study amphioxus-specific patterns. We only considered protein-coding genes, where orthologs and paralogs can be more easily identified using sequence similarity (see Methods for protein-coding gene set filtering). To detect orthogroups we used Broccoli [32], an algorithm based on protein sequence similarity designed to split gene trees with duplications older than the first speciation event in the tree into different orthogroups. That is, with our set of species, this algorithm classifies gene duplications predating chordates into different orthogroups and, thus, they are not counted as duplications in our study. In other words, we only consider duplications posterior to the split of vertebrate and amphioxus lineages (Figure 2A).

**Figure 2.**
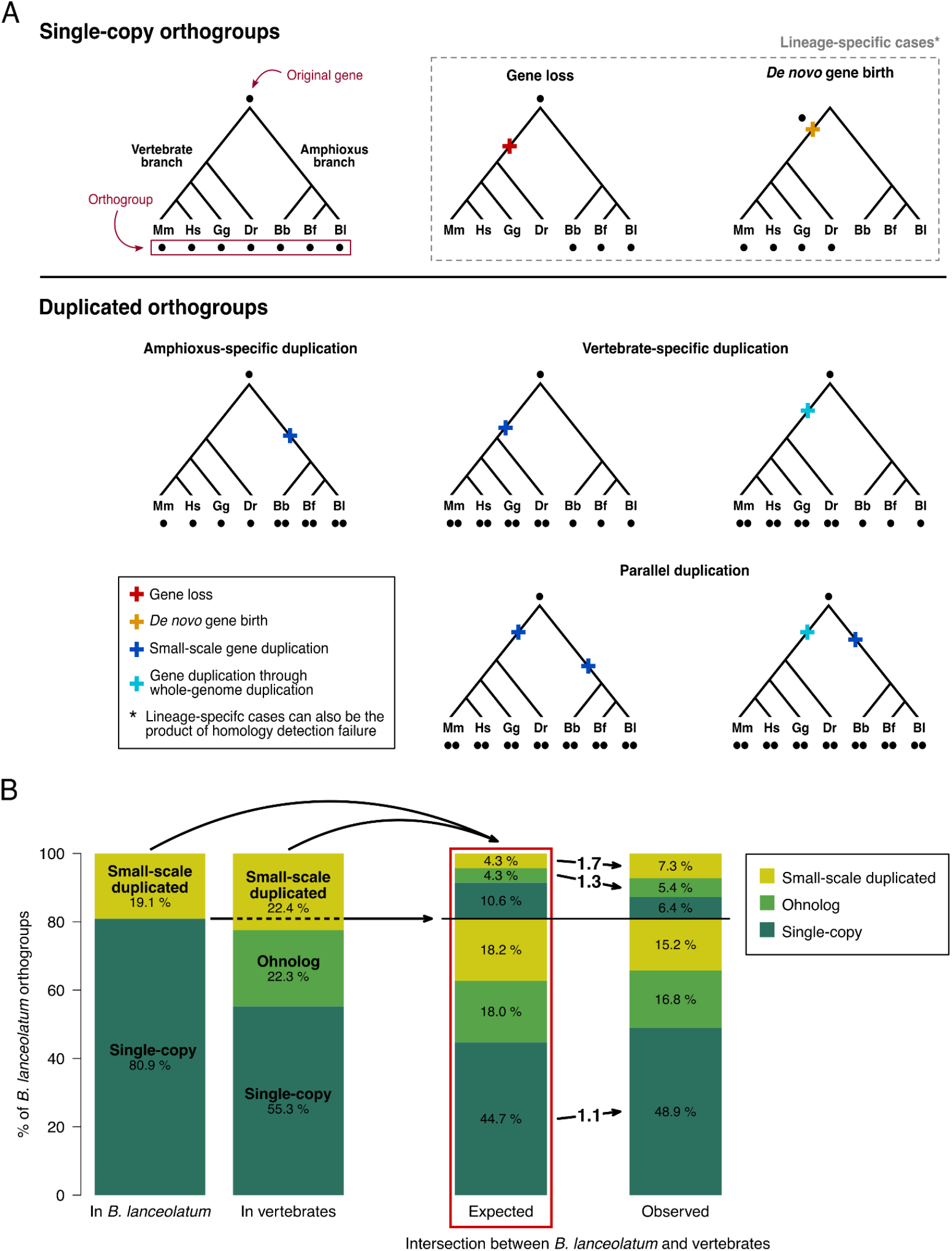
Single-copy and duplicated orthogroups in the amphioxus and vertebrate lineages. **A.** Schematic representation of the chordate gene orthogroup classification, showing examples of different gene evolutionary histories represented as gene trees. Each point represents a gene, all gene trees are derived from a single original gene and all genes at the tips of a tree are grouped into an orthogroup. Orthogroups are classified as duplicated or single-copy depending on whether there was a duplication in their evolutionary history or not. Duplicated orthogroups can originate from amphioxus-specific, vertebrate-specific, or parallel duplications. In addition, gene duplications in the vertebrate lineage can be the product of either a small-scale duplication or a whole genome duplication (ohnologs; see Methods). All duplications in the amphioxus lineage are small-scale. Duplications predating the last common ancestor of chordates are separated in different orthogroups and, thus, not taken into account as duplications in our study (see Methods). Events of gene loss, *de novo* gene birth or homology detection failure are also considered. These events can result in lineage-specific orthogroups and can appear in any branch of the tree and in combination with duplication events. Each orthogroup gene evolutionary history is reconstructed based on the number and distribution of its genes across vertebrate and amphioxus species. Mm, Hs, Gg, Dr, Bb, Bf and Bl correspond to mouse (*M. musculus*), human (*H. sapiens*), chicken (*G. gallus*), zebrafish (*D. rerio*), and *B. belcheri, B. floridae* and the European amphioxus (*B. lanceolatum*), respectively. **B.** Enrichment in parallel duplications in the amphioxus and the vertebrate lineages. Only *B. lanceolatum* orthogroups which have vertebrate orthologs are represented. The 1st column on the left shows the percentage of *B. lanceolatum* orthogroups that are either single-copy or small-scale duplicated. The 2nd column shows the percentage of *B. lanceolatum* orthogroups that are either single-copy, ohnolog or small-scale duplicated in vertebrates. The 3rd and 4th columns show, respectively, the expected (if independently distributed) and the observed intersections between the sets of orthogroups depicted in columns 1 and 2. In the 3rd and the 4th columns, orthogroups below the horizontal line correspond to orthogroups that are single-copy in *B. lanceolatum* while orthogroups above the horizontal line correspond to orthogroups duplicated in *B. lanceolatum*. Categories that are enriched in the observed values, compared to the expected, are depicted with arrows between the 3rd and the 4th columns. Numbers on those arrows correspond to the binary logarithm of the ratio between observed and expected values (log2 fold change), all of which have p-value ≤ 0.0002 after Bonferroni multiple testing correction (corresponding values are in Table S1). For example, 4.3% of the *B. lanceolatum* small-scale duplicated orthogroups are expected to be small-scale duplicated in vertebrates (top light green set in the 3rd column) while 7.3% is observed (top light green set in the 4th column). This represents a 1.7 log2 fold change enrichment.

Among all chordate orthogroups, we distinguished between single-copy orthogroups, lineage-specific duplicated orthogroups (either amphioxus- or vertebrate-specific), parallel duplicated orthogroups (duplicated independently in both lineages), and lineage-specific orthogroups (Figure 2A). This last type of orthogroups (lineage-specific orthogroups) are either born *de novo* in the lineage where they are present, are the result of a gene loss in the other lineage, or have diverged enough to elude sequence-similarity based orthology (homology detection failure). For duplications in vertebrates, we distinguish between *small-scale duplicated genes* and *ohnologs*, the latter corresponding to genes duplicated during the 2R rounds of genome duplication at the origin of vertebrates. In order to retrieve patterns common to all vertebrates, teleost-specific ohnolog genes were not considered (see Methods for specifications on ohnolog definition). In the case of amphioxus, all gene duplications were considered small-scale duplications. To make sure that we were not considering duplications older than the split between amphioxus and vertebrates, we validated Broccoli results with an independent phylogenetic approach. We were able to reconstruct phylogenetic trees using RAxML for 69.14% of orthogroups that are small-scale duplicates both in vertebrates and in amphioxus according to Broccoli. Of these, 79.95% were validated as having all vertebrate and all amphioxus genes in two different monophyletic groups. Similar results were obtained when considering orthogroups which have ohnologs in vertebrates and small-scale duplicates in amphioxus (66.37% of reconstructed trees and 74.83% of monophyletic grouping of both vertebrate and amphioxus genes).

We detected 8705 orthogroups shared by amphioxus and vertebrates (65.6% of amphioxus groups and 68.4% of vertebrate groups). Thus, a large part of gene orthogroups are common and still detectable by sequence similarity, despite 500 million years of independent evolution between the two lineages. This is the set of shared orthogroups between amphioxus and vertebrates that we have considered, although others are likely to exist which are more divergent in sequence and thus undetected with the methods used (homology detection failure) [33]. We identified 4559 amphioxus lineage-specific orthogroups. As stated above, these orthogroups are either lost in vertebrates, born *de novo* in the amphioxus lineage, or a product of homology detection failure. The majority of these amphioxus-specific orthogroups are shared among the three amphioxus species included in the analysis (Figure S2A). *B. lanceolatum* shares slightly more orthogroups with *B. floridae* than it does with *B. belcheri,* consistent with the known *Branchiostoma* phylogeny in which *B. lanceolatum* and *B.floridae* are sister groups [34,35]. *B.floridae* and *B. belcheri* share less orthogroups than each of them does with *B. lanceolatum,* possibly due to differences in genome assembly and annotation, which supports the higher quality of BraLan3.

Around half of all genes present in each species are duplicated, in both vertebrate and amphioxus genomes (Table 2). These results suggest that duplication contributes to the amphioxus gene repertoire in similar proportions to that of vertebrates. On the other hand, they do differ in the distribution of duplications among orthogroups. Slightly less than 20% of amphioxus orthogroups have duplications, while consistently more than 20% of orthogroups in vertebrates do. We have purposely avoided quantifying 3R zebrafish ohnologs as ohnolog orthogroups in order to focus on 2R ohnologs. Despite this measure, we do see a higher percentage of duplicated genes in the zebrafish genome because zebrafish 3R ohnolog orthogroups were included if there was another duplication in one of the other vertebrate species. The average number of duplicated genes in orthogroups with duplications is consistently higher in amphioxus than in vertebrates. The examination of the chromosomal location of duplicated genes showed that tandemly duplicated genes are more abundant in *B. lanceolatum* than in vertebrates (Figure S2B). In conclusion, amphioxus and vertebrate genomes have similar proportions of duplicated genes but their distribution across orthologous groups is different, very likely because of the existing differences in gene duplication mechanisms between the two lineages, namely, the existence of two rounds of the 2R rounds of WGD in vertebrates and no WGD in the amphioxus lineage [6,36].

**Table 2.** Gene duplication prevalence in different species of amphioxus and vertebrates. Genes in each species were filtered for proper correspondence between gene annotation and protein sequence (see numbers in Table S6). For example, from the 26,468 *B. lanceolatum* genes with strong evidence (see Methods) we used 25,946 genes for the orthology analysis. Percentages are calculated from the total number of genes or the total number of orthogroups in each species, respectively.

| Lineage | Species | Genes |  |  | Amphioxus - Vertebrate orthologous genes |  |  | Orthogroups |  |  | Mean # of genes in duplicated orthogroups |
| --- | --- | --- | --- | --- | --- | --- | --- | --- | --- | --- | --- |
|  |  | Total | Duplicated | % | Total | Duplicated | % | Total | Duplicated | % |  |
| Amphioxus | *B. lanceolatum* | 25946 | 11973 | 46.1 | 15269 | 8520 | 55.8 | 12691 | 2468 | 19.4 | 4.9 |
|  | *B. floridae* | 20541 | 10192 | 49.6 | 13074 | 7578 | 58.0 | 10568 | 1934 | 18.3 | 5.3 |
|  | *B. belcheri* | 18413 | 8469 | 46.0 | 11944 | 6354 | 53.2 | 10533 | 1805 | 17.1 | 4.7 |
| Vertebrates | *D. rerio* | 23912 | 13510 | 56.5 | 16664 | 11504 | 69.0 | 10587 | 3727 | 35.2 | 3.6 |
|  | *G. gallus* | 16558 | 7416 | 44.8 | 12250 | 6630 | 54.1 | 10455 | 2399 | 22.9 | 3.1 |
|  | *M. musculus* | 21691 | 10660 | 49.1 | 14447 | 8518 | 59.0 | 12236 | 2847 | 23.3 | 3.8 |
|  | *H. sapiens* | 19715 | 9799 | 49.7 | 13802 | 8015 | 58.1 | 12205 | 2997 | 24.6 | 3.3 |

Gene duplication patterns are conserved between amphioxus and vertebrates: genes which are duplicated in one lineage tend to be duplicated in the other lineage (Figure 2B, Table S1). *B. lanceolatum* small-scale duplicated orthogroups are enriched (1.7-fold enrichment) in small-scale duplicated vertebrate orthogroups (Figure 2B). Surprisingly, they are also enriched in vertebrate ohnologs (1.3-fold enrichment; Figure 2B). That is, what is duplicated in one lineage tends to be duplicated in the other lineage regardless of the gene duplication mechanism, either through small-scale duplication or WGD (ohnologs). There is also an enrichment in orthogroups which are single-copy in both lineages (1.1-fold enrichment; Figure 2B). All these results are statistically significant (p-value ≤ 0.0002 after Bonferroni multiple testing correction; values are in Table S1), are robust to the inclusion of lineage-specific genes in the analysis (Table S2), and are consistent with recent observations in other amphioxus species [11]. Moreover, duplicated orthogroups with a low number of gene copies (<=2.5 mean number of genes) in amphioxus tend to have a low number of gene copies in vertebrates (Spearman’s *ϱ* = 0.28; Table S3, contingency table chi-squared test p-value < 0.0001). This is true for both small-scale duplicates and ohnologs (Table S3).

### Functional patterns of duplicate genes

We wanted to know whether the parallelism in duplication between vertebrate and amphioxus lineage shown in the previous section extends to the level of functional categories. In order to group vertebrate and *B. lanceolatum* genes in functionally related groups, we have used the Gene Ontology (GO) terms that annotate genes to hierarchically organized functional categories [37,38]. We used both *molecular function* GO terms and *biological process* GO terms, which refer respectively to molecular-level activities performed by gene products, and to larger processes accomplished by multiple molecular activities [37,38].

There is a strong positive correlation in the proportion of duplications in different functional categories between the vertebrate and the amphioxus lineages (Figure 3A for *molecular function* GO terms and 3B for *biological process* GO terms). The categories with the lowest rates of duplication in both lineages correspond to basic functional categories, such as mitochondrial, transcriptional, translational or cell cycle-related *biological process* GO terms or ribosome, DNA, RNA and nuclear-related *molecular function* GO terms (Table S4). Conversely, many GO terms related to regulation, signaling, immune system or response have more than 80% or even 90% of genes duplicated in both lineages (see Table S4 for details). These results suggest strong selection for preserving single-copy genes in some functions, and either a tolerance to duplication or selection favoring duplication in others. Of special interest is that the strength of the trend is much larger when both types of vertebrate duplicates (ohnologs and small-scale duplicates) are taken into account together instead of separately (Figures 3A for *molecular function* GO terms and 3B for *biological process* GO terms). This means that amphioxus small-scale duplications alone share most of the functional constraints on (or tolerance to) duplication that affect both ohnologs and small-scale duplications in the vertebrate lineage. Together, these results highlight that, despite well established results on different constraints on paralog retention between small-scale duplicates and ohnologs [39–42], selection for preserving single-copy genes in certain basic functional categories drives a strong signal of parallelism in copy number between sub-phyla.

**Figure 3.**
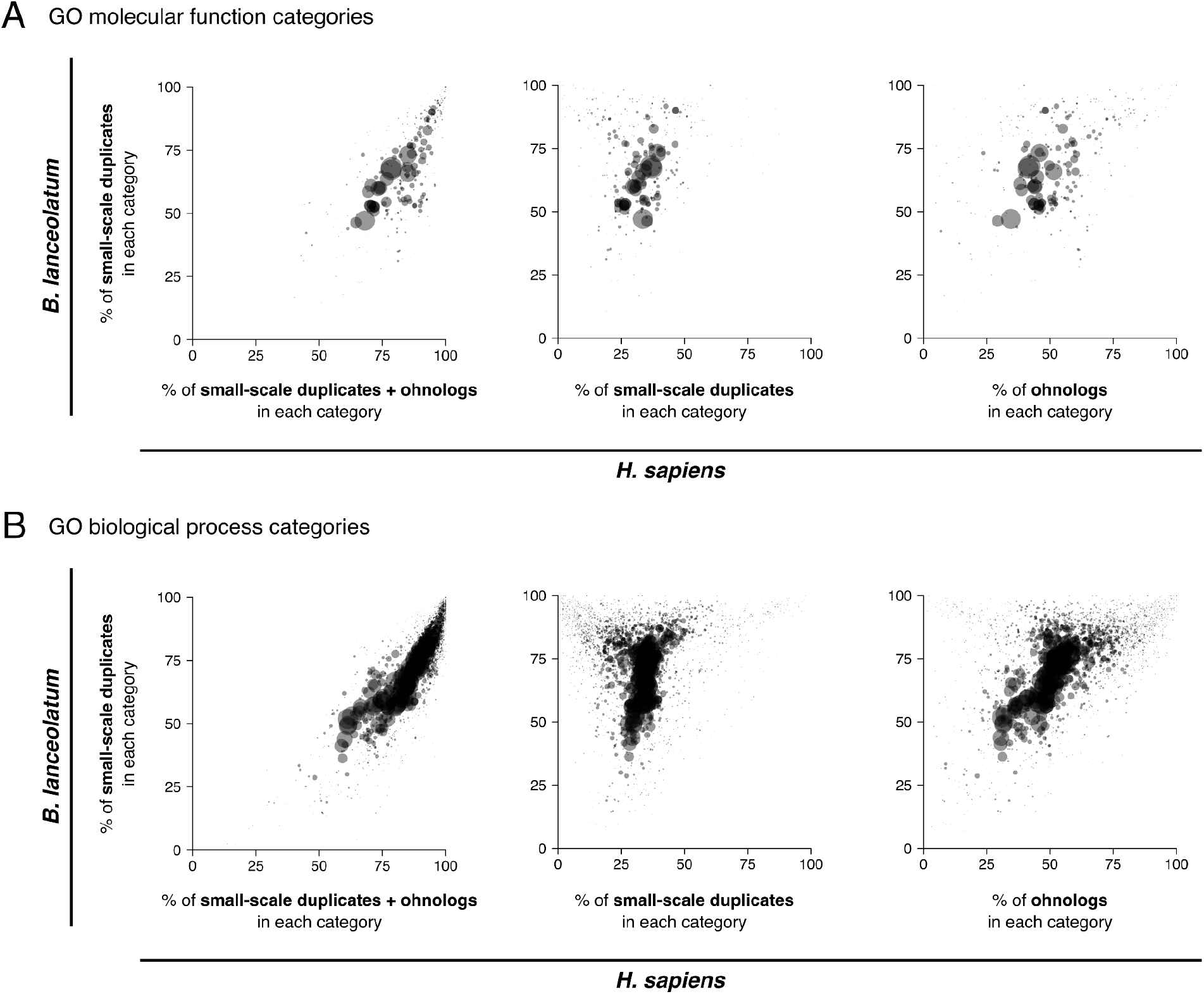
Parallelism between amphioxus and human in the amount of gene duplicates across Gene Ontology (GO) terms. Each point in each plot represents a GO term, where point size is proportional to the number of *H. sapiens* genes in the corresponding GO term. The percentage of the total number of genes annotated to each GO term which are small-scale duplicates in *B. lanceolatum* (y-axis in all plots) is compared with the percentage of the total number of genes annotated to the same GO term that are duplicated in *H. sapiens* (x-axis), including both small-scale duplicates and ohnologs, (left); only small-scale duplicates (center); and only ohnologs (right). *Molecular function* GO terms are represented in **A** and *biological process* GO terms in **B**. For example, 35.5% of the 1551 human genes annotated to the biological process GO term “cell cycle” (GO:0007049) are duplicated by small-scale duplication, while 56.1% of the 1077 amphioxus genes annotated to the same term are duplicated; thus this term is represented at the coordinated (56.1, 35.5) in the graph in the middle of **B**. *B. lanceolatum* GO term annotation was extrapolated from the human GO term annotation, and only *H. sapiens - B. lanceolatum* orthologous genes were considered in this analysis (see Methods). Only GO terms with a minimum of 50 genes in both human and *B. lanceolatum* were considered. No statistical test for correlation was performed since GO terms are not independent of each other. All points are filled with the same semi-transparent black color, thus opaque regions imply a high point density.

### Expression and evolution of duplicate genes

To characterize *B. lanceolatum* duplicated genes independently of GO annotations in human, we re-analysed *B. lanceolatum* RNA-seq from Marlétaz et al. [8]. Amphioxus duplicated genes have lower levels of expression than single-copy genes. This is true both in adult tissues (Figure 4A) and during embryonic development (Figure S3A), and whatever the profile of the corresponding vertebrate orthologs: single-copy, ohnologs, small-scale duplicates, or *B. lanceolatum*-specific (Figure 4B and Figure S3B). The lower expression of amphioxus duplicated genes with respect to single-copy genes is preserved within functional categories (Figure S4). Furthermore, amphioxus duplicated genes show higher levels of adult tissue-specificity and developmental stage-specificity than single-copy genes (Figure 4C and Figure S3C). Again, this is true independently of the profile of the corresponding vertebrate orthologs (Figure 4D and Figure S3D). Regarding *B. lanceolatum*-specific genes, they show lower expression and higher levels of tissue and developmental stage specificity compared to genes with orthologs, even when they are single-copy (Figures 4B and Figure S3B). This is an expected result given that this category contains rapidly-evolving (including homology detection failures) and young genes.

**Figure 4.**
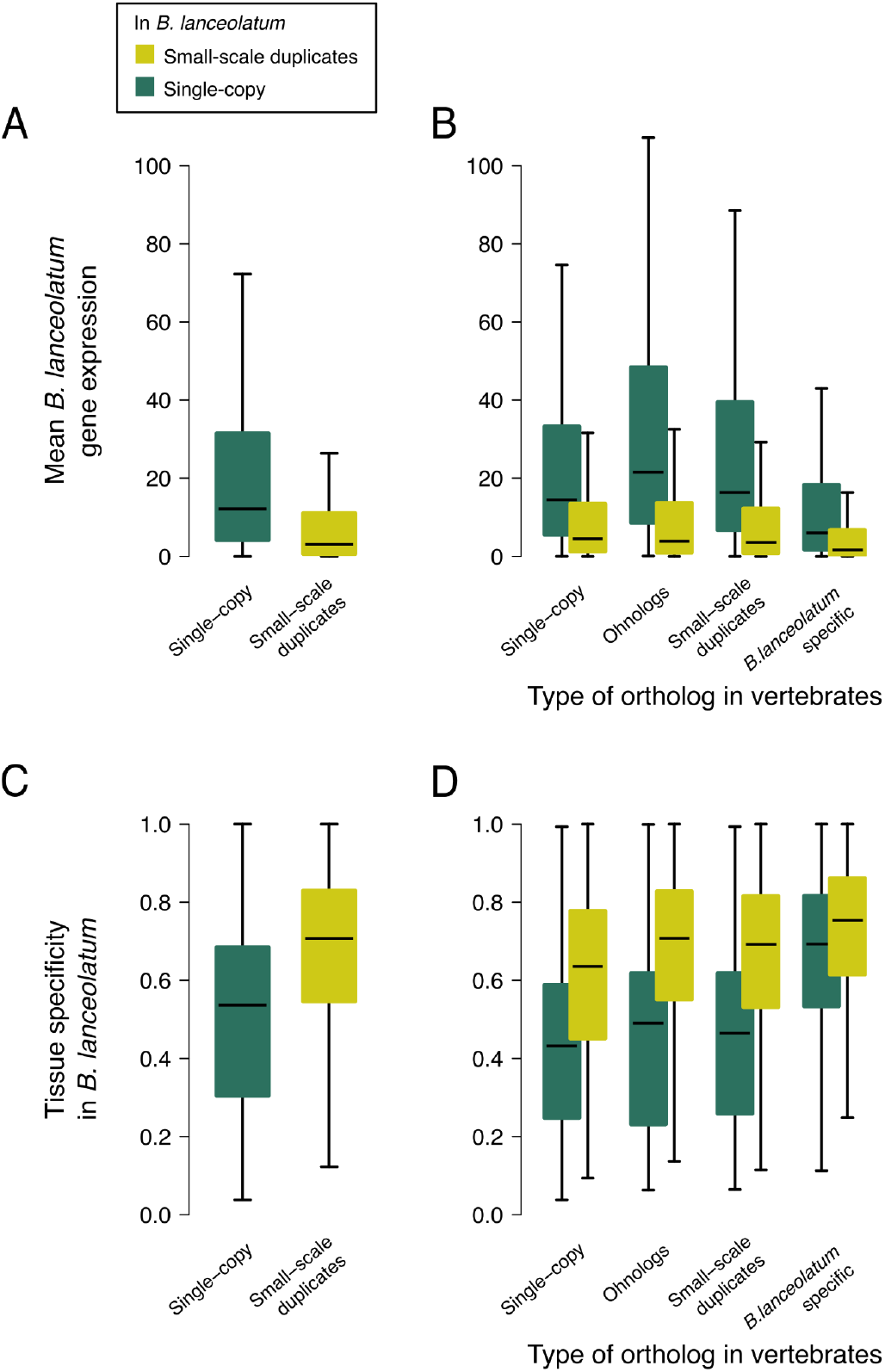
Gene expression in single-copy and duplicated genes in *B. lanceolatum*. **A.** Box plots showing the distribution of mean gene expression (in transcripts per million; TPM) across adult amphioxus tissues for *B. lanceolatum* single-copy and small-scale duplicated genes. Outliers are not represented (see Methods). **B.** Same as in **A** but dividing *B. lanceolatum* genes according to the status of their vertebrate ortholog if present (single-copy, ohnolog, small-scale duplicate, or no ortholog [*B. lanceolatum* specific]). **C-D**. Box plots showing tissue specificity distribution of *B. lanceolatum* gene expression (Tau statistic) for the same gene groups as **A** and **B**, respectively. Tau values range from 0 (ubiquitous expression) to 1 (expression only in 1 tissue). Similar results across developmental stages are shown in Figure S3.

Together, these results show, on the one hand, that amphioxus single-copy genes have a major role in the transcriptome of amphioxus. They show higher gene expression and less tissue and developmental stage specificity, indicating a role in the maintenance of basic cellular processes in amphioxus. This trend is independent of their vertebrate orthologs’ duplication profile, suggesting that gene expression is more dependent on the specific duplication status in amphioxus rather than on the long term evolutionary constraints in duplicability. On the other hand, amphioxus duplicates have lower expression and higher specificity than single-copy genes, suggesting secondary but more specific roles of duplicated genes in this species. These results are consistent with the analysis of functional categories, where duplicated genes are less present in core functional categories and less expressed whenever they are present, and coherent with previously described patterns of expression of duplicated and young genes [43–48].

In order to further explore the evolution of amphioxus duplicated genes, we compared expression in amphioxus and zebrafish orthologs (following the approach of Marlétaz et al. [8]). Presence or absence of gene expression in 7 *B. lanceolatum* conditions (embryo, male gonads, muscle, neural tube, gut, gills and hepatic diverticulum) was matched to the presence or absence of gene expression in 7 homologous zebrafish conditions (embryo, testis, muscle tissue, brain, intestine, pharyngeal gill and liver, respectively). Gene expression data was retrieved from Marlétaz et al. [8]. For every pair of orthologs, the difference in the number of expressed conditions between *B. lanceolatum* and zebrafish was calculated (number of expressed *B. lanceolatum* conditions minus number of expressed *D. rerio* conditions). The distribution of these differences for orthologous pairs of genes that are single-copy in both species is symmetric, with a strong centrality around 0 showing an overall conservation of expression in homologous conditions between the two species (Figure 5, first case). We observe a loss in the number of expression conditions of vertebrate ohnolog genes compared to single-copy orthologs in amphioxus (Figure 5, second case; black arrow), replicating results by Marlétaz et al. [8]. We also confirm their result that this tendency (skewness) is reverted when taking both duplicates’ gene expression conditions into account (union of duplicates; brown line in Figure 5). These results suggest either specialization or subfunctionalization of vertebrate ohnolog genes in different gene expression conditions with respect to the single-copy amphioxus ortholog. Here, we show that this has also happened both in vertebrate small-scale duplicates (third case in Figure 5) and in amphioxus small-scale duplicates (fourth case in Figure 5). That is, for all types of gene duplications (ohnologs and small-scale duplicates) and in both the vertebrate and amphioxus lineages, we observe specialization or subfunctionalization of gene duplicates in the number of conditions of expression. Moreover, in all cases, when all duplicated genes in an orthogroup are accounted for together, they tend to have an expression profile similar to the one of their single-copy ortholog (reversion of skewness in brown lines in Figure 5). We also observe the same pattern when splitting amphioxus duplicated genes between inter-chromosomal, distant intra-chromosomal, and tandemly duplicated genes (Figure S2C). These results support a general trend in all types of duplicates in both lineages that is consistent with expectations for subfunctionalization or specialization.

**Figure 5.**
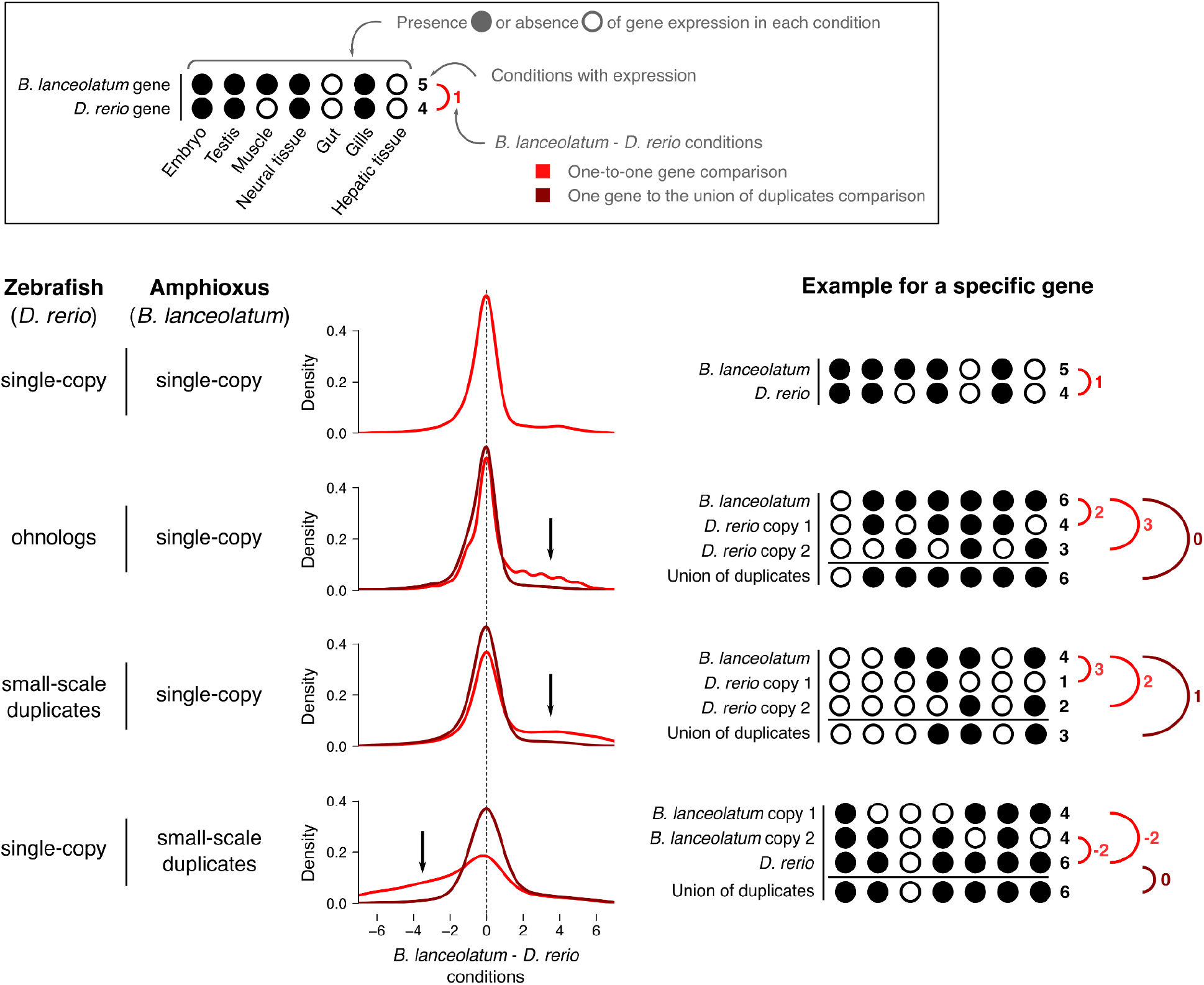
Specialization of duplicated genes in amphioxus and zebrafish. Amphioxus average gene expression in embryonic stages, male gonads, muscle, neural tube, gut, gills and hepatic diverticulum was matched to zebrafish gene expression in embryo, testis, muscle tissue, brain, intestine, pharyngeal gill and liver, respectively. The presence or absence of gene expression in each condition (either adult tissue or developmental stage whole body) and each species was determined, and the difference in number of expressed conditions between the two species was calculated (*B. lanceolatum* - *D. rerio* conditions) for every pair of orthologous genes (one-to-one gene comparison, in red). In the case of duplicated genes, the *union of both duplicates’ expression* was calculated as the number of conditions in which either of the duplicates is expressed (see examples in figure). The union of duplicates was then compared to their ortholog(s) gene expression in the other species (comparing one single gene to the union of duplicates, in brown). The density of the distribution of this difference in expression between *B. lanceolatum* and *D. rerio* is shown for four different types of orthologous genes: genes that are single-copy in both species (first row); zebrafish ohnolog genes that are orthologs to *B. lanceolatum* single-copy genes (second row); zebrafish small-scale duplicates that are orthologs to single-copy *B. lanceolatum* genes (third row); and zebrafish single-copy genes that are orthologs to small-scale duplicates *B. lanceolatum* genes (fourth row). Black arrows highlight the direction of the skewness (when present) of the one-to-one gene comparisons (red lines). Hypothetical examples of each case are schematically shown on the right in order to make the figure clearer.

### Amphioxus gene synteny

*B. lanceolatum* and *B.floridae* divergence time estimates range from 20 to 150 million years ago (Mya), while the amphioxus lineage diverged from that of vertebrates and tunicates around 500 Mya [34,35]. *B. lanceolatum* and *B. floridae* shared their evolution for most of the time since the amphioxus lineage split with that of other chordates. The presence of chromosome scale genome references for these two species [9] allows for comparison of gene order. While one-to-one orthologs show an overall conservation of synteny between the two species, there are also two important chromosomal rearrangements (Figure 6). Although both species share the same number of chromosomes (n = 19), we observe two large chromosomal fusions and fissions between the two species. The largest chromosome of *B. lanceolatum* (chromosome 1) appears to be homologous to two smaller *B. floridae* chromosomes (chromosomes 9 and 11). Conversely, chromosome 3 of *B. floridae* appears to be homologous to *B. lanceolatum* chromosomes 15 and 19. In order to shed light on the ancestral state of these chromosomal fusions or fissions, we compared the *B. lanceolatum* and *B. floridae* synteny to the fragmented *B. belcheri* genome reference at the gene level (Figure S5). We found no evidence for either chromosomal fusion in *B. belcheri.* That is, none of the *B. belcheri* scaffolds map to both chromosomes 9 and 11 of *B.floridae* or to both chromosomes 15 and 19 of *B. lanceolatum.* These results match recent evidence on *B. belcheri* and *B.floridae* karyotype reconstruction, pointing towards a chromosomal fusion in the *B. floridae* branch with respect to *B. belcheri* [11]. Here, we show that this chromosomal fusion is *B. floridae* specific (is not shared with *B. lanceolatum*) and that there is an additional chromosomal fusion in *B. lanceolatum*.

**Figure 6.**
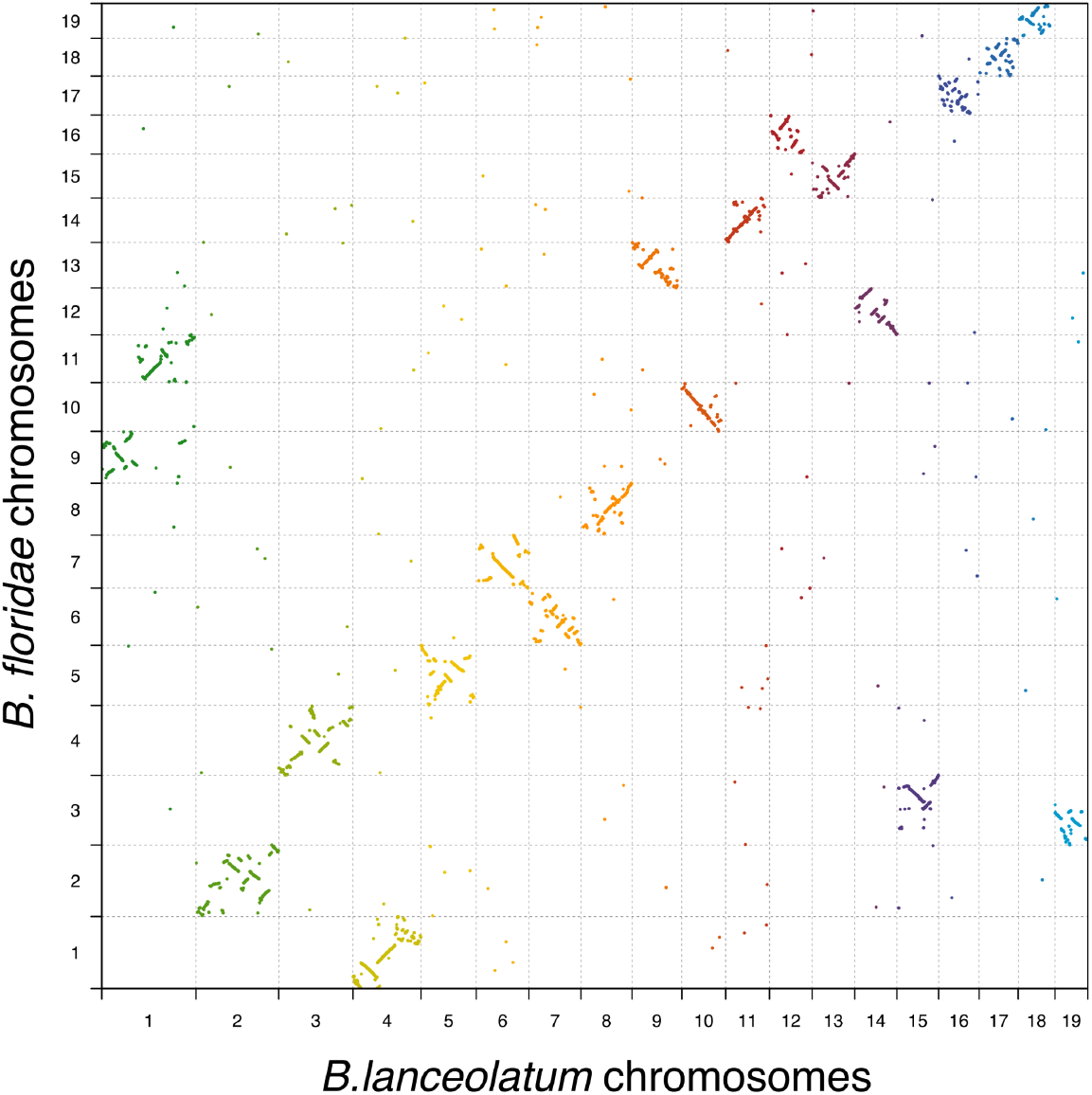
Single-copy genes synteny conservation between *B. lanceolatum* and *B. floridae*. Every point represents a gene that is single-copy in both species, and it is represented by its midpoint coordinates in each genome. Different colors depict different *B. lanceolatum* chromosomes.

Gene collinearity of one-to-one orthologs between *B. lanceolatum* and *B. floridae* is generally conserved, with frequent medium scale intra-chromosomal rearrangements (inversions and translocations). This observation is true for the majority of chromosomes but fails for the smallest chromosomes (*B. lanceolatum* chromosomes 16, 17 and 18 that correspond to *B.floridae* chromosomes 17, 18 and 19 respectively) where we find a loss of gene collinearity and a dispersed distribution of one-to-one orthologs. Unlike one-to-one orthologs, gene duplicates show a scattered pattern of chromosome distribution between *B. lanceolatum* and *B.floridae* (Figure S6A), with local gene expansions. As expected, they show no evidence of any large-scale duplication event and show that both intra- and inter-chromosomal small-scale duplications have been frequent in the amphioxus lineage. This is confirmed in the *B. lanceolatum* intraspecies duplicated gene synteny where we observe frequent intra- and inter-chromosomal gene duplications (Figure S6B). Local expansions of genes are frequently found among *B. lanceolatum* paralogs, but are rarer among paralogs between *B. lanceolatum* and *B. floridae*, suggesting a dynamic paralog synteny.

Between *B. lanceolatum* and vertebrate (*G. gallus* in Figure S6C) one-to-one orthologs, we observe several conserved synteny blocks with a complete loss of gene collinearity. This is consistent with previously reported results for *B. floridae* [9,10], and suggests that in addition to chromosomal rearrangements, there were numerous middle and small-scale rearrangements affecting genes in both lineages, as commonly observed in vertebrates [21,49]. These have broken gene collinearity while preserving the context of conserved synteny blocks. Interestingly, *B. lanceolatum* chromosomes 10, 12 and 17 share a very similar pattern of gene synteny with *G. gallus* (Figure S6C) suggesting a shared chromosomal history of these three amphioxus chromosomes. This pattern is supported by similar observations for three *B. floridae* chromosomes [9]. In the case of duplicated genes, we find some synteny block conservation between chordate lineages with a general dispersal of duplicated genes across the genome. We also observe multiple local gene expansions, more frequent in the vertebrate lineage than in the amphioxus one (Figure S6D).

## Discussion

Amphioxus are generally characterized by slow molecular evolution, notably slow protein sequence evolution, relative to the two other lineages of chordates, ascidians and vertebrates [6,7]. This provides a useful contrast to the vertebrate diversity of phenotypes and genomes, and has led to the use of amphioxus to understand many features of vertebrate evolution, serving as a proxy for the ancestral state. Given the role of gene duplication in evolutionary change, one could expect to find a low duplication rate and a minor role of duplicate genes in the amphioxus lineage. Yet, we and others [11] find a similar amount of small-scale gene duplication activity in amphioxus as in vertebrates, and we show a strong parallelism in terms of genes, biological functions, genome distribution, and gene expression profiles.

The extent of amphioxus-lineage duplication has been difficult to study until recently. This is because the high heterozygosity of amphioxus genomes was a problem for the reliable assembly of duplicated genes [6,24]. In this study, we constructed both high-quality assembly and annotation for *B. lanceolatum*, allowing us to infer a reliable set of paralogs, and to study these duplications genome-wide. The usage of long-read and long-range scaffolding data as well as specifically designed methods for alternative haplotype filtering provided a high quality reference on top of which we cautiously annotated genes. As is usual when annotating the genome of a non-model organism, we filtered gene models by similarity to known genes or proteins. This can lead to false negatives, especially putative faster-evolving duplicate or lineage-specific genes. To minimize this issue while avoiding false positives, we also retained both gene predictions with a BLAST hit in other amphioxus annotations (recovering 143 genes) and gene predictions expressed above the background in at least three samples (recovering 986 genes). This has allowed us to recover a total of 11973 duplicated genes in *B. lanceolatum*, a similar proportion of the total gene set as in most vertebrates. With different methods, Huang et al. [11] recently reported high quality genomes for several other species of amphioxus and also found similar proportions of gene gains (including duplicated genes) in the vertebrate and amphioxus lineages.

The parallelism of duplication patterns between vertebrates and amphioxus is striking. The same functional categories are enriched or depleted. Notably, regulatory genes are over-represented in amphioxus duplicated genes, as they are in other lineages, including yeasts, plants and vertebrates [50,51]. This shows that amphioxus have continued to evolve complex regulatory processes or pathways, despite remaining apparently simpler than other chordates. This is reinforced by the parallelism of duplicate gene evolutionary patterns, with a similar dominance of expression specialization or subfunctionalization in amphioxus and vertebrates. Moreover, synteny analysis shows a dispersed distribution across the genome of small-scale duplicates in the amphioxus lineage, similar to the one known for vertebrates [52], pointing towards common gene duplication mechanisms and dynamics. Of note, Huang et al. [11] report an excess of segmental duplications in other amphioxus species.

While our new genome assembly confirms the overall conservative evolution of amphioxus synteny, we find chromosomal rearrangements between *B. lanceolatum* and *B. floridae*. Our observations corroborate the conclusions derived from the very recently reported high quality genomes of three other amphioxus [11] that the ancestral *Branchiostoma* karyotype had 20 pairs of chromosomes, and we report two independent chromosomal fusions in *B. lanceolatum* and *B. floridae*. These rearrangements, together with the duplication patterns, illustrate how amphioxus genomes continue to be shaped by large scale evolutionary events.

## Conclusions

The amphioxus lineage has a history of small-scale gene duplications similar to the one observed in the vertebrate lineage, and there is a conservation of the constraints on gene duplication between these two lineages. Our results highlight the durability of the selection preventing duplication in certain gene families and allowing or favoring it in others. In the amphioxus and vertebrate lineages, around 500My of independent evolution and the large diversification of vertebrate’s phenotypes have not erased most of these constraints on gene duplication despite the 2R WGDs in early vertebrate evolution.

## Methods

### Genome assembly

We generated 9.46M PacBio reads through 20 cells of PacBio RSII, with a N50 of 10998 and totalling 80.2Gb of raw data, which represents 146x coverage of the haploid *B. lanceolatum* genome. Canu (v1.9) was run on the PacBio data using the parameters correctedErrorRate=0.065, ovlMerDistinct=0.975 and batOptions=“-dg 3 -db 3 -dr 1 -ca 500 -cp 50” to maximize haplotype separation [53]. With these settings, Canu yielded, a diploid assembly of 931.9Mb with a N50 of 1.02Mb that was subjected to polishing with the Arrow algorithm (v2.3.2) as implemented in the GenomicConsensus package (PacBio) after aligning back the raw PacBio reads with Pbmm2 (v1.1.0) relying on Minimap2 (v2.17). After polishing, we filtered haplotigs and heterozygous regions from the assembly with purge_dups (v1.0.0), relying on coverage depth and sequence reciprocal alignment [54]. We estimated coverage by aligning PacBio reads with Minimap2, and we used a coverage cutoff of 105 to distinguish between homozygous and heterozygous regions. We further scaffolded the resulting haploid assembly (size: 507.4Mb with a N50 of 1.48Mb) using chromatin conformation capture data (HiC data). HiC data was aligned and duplicated and spurious read pairs were filtered out using Juicer (v6be7c0f) [55] and BWA-MEM (v0.7.17). The subsequent read pairs were used as an input by 3D-DNA to perform contact-based scaffolding [56]. The resulting assembly was manually edited using Juicebox to correct problems and subjected to a final round of optimization with 3D-DNA. This assembly notably had 19 large scaffolds consistent with the chromosome number of *B. lanceolatum* [29], in addition to multiple smaller scaffolds. To reduce the computational load of downstream analyses, we removed small scaffolds that did not have multi-intronic gene models or that had a high percent (≥50%) of repeat content.

Two additional modifications were done to this intermediate assembly. First, the assembly of the Irx cluster on chromosome 7 was manually curated based on previous information [57]. In short, the ortholog of *IrxC* was already located on chr7, but *IrxA* and *IrxB* were each present in a different small scaffold, which were introduced in the correct location of chr7. Second, we detected various gene models that contained in-frame stop codons. Upon further inspection, these were found to be due to indels introduced by PacBio sequencing. To correct these errors, we extracted all unique exons obtained through an initial gene annotation with Mikado and StringTie+Transdecoder (see below) and blasted them against a combined Trinity assembly generated with Illumina RNA-seq data from multiple tissues and developmental stages [8]. The blast output was parsed to conservatively detect single indels within 10 nt alignment windows, supported by at least five different Trinity transcripts coming from at least two independent RNA-seq samples. These indels corresponded to 3672 insertions and 2690 deletions with respect to the Trinity assembly, mainly in UTR exons and/or lowly supported gene models, and were edited in the genome sequence to produce the final assembly.

### Annotation

Stage- and organ-specific transcriptomes (RNA-seq) [8] were aligned using STAR and assembled as individual transcriptomes using StingTie [58]. The StingTie assemblies (GTF files) were merged using Taco [59]. The RNA-seq data was also assembled in bulk with Trinity as reported in Marletaz et al. 2018 [8] and aligned to the new assembly using Minimap2 with the ‘splice:hq’ parameter. These transcriptomes were leveraged using the Mikado tool [60]. We also generated spliced protein alignment using Exonerate from the annotation of the *B. floridae* genome [9] assuming at least 65% protein identity and a maximum intron size of 250kb. We converted the Mikado transcriptome assembly and the Exonerate alignment into hints for the Augustus gene prediction tool [61], as ‘exon’ and ‘CDS’ hints, respectively. Augustus was run using the previously defined model and the aforementioned hints while allowing hinted splices (‘ATAC’). Augustus yielded 37,787 gene models, of which 7,101 overlapped with a repeat (all exons with at least 50% overlap with repeats) and were excluded. We constructed a repeat library using RepeatModeller (v2.0) and masked repeats using RepeatMasker (v4.0.7). We also generated a transcriptome dataset by aligning the assembled transcriptome using PASA (v2.3.2) and used it to add UTRs and isoforms to the Augustus gene models through two rounds of processing.

A series of additional corrections were then applied to this initial annotation. First, 5’ and 3’ UTRs were extended based on a GTF merge of the StingTie assemblies on which Transdecoder (PMID: 23845962) was run to identify open reading frames. 3’ UTRs were extended up to 5 Kbp unless they overlap with downstream models in the same strand and provided these 3’ UTRs had read support (otherwise, only 0.5 Kbp extensions were allowed). Upon extension, the end of the gene models were modified if there was no annotated STOP codon and an in-frame STOP codon was added with the UTR extension. Similarly, for gene models with no annotated start codon, 5’ UTRs were extended when possible based on StingTie information, and a start codon were added when an upstream in-frame ATG was introduced with the extension. In addition, we noticed some Augustus gene models whose annotated start codon seemed upstream than expected based on a protein alignment with the human orthologs. To identify and correct these cases, when the initial M of the human ortholog aligned with an M in the amphioxus sequence, this M was selected as the new starting codon. Finally, we performed a search for potential chimeras and broken genes comparing Augustus (default) and StingTie+Transdecoder gene models, as described in [8]. These pairs of overlapping models were manually inspected and corrected by substituting the Augustus gene models by StingTie+Transdecoder ones when appropriate. Finally, gene models were named based on the following nomenclature, BLAGXXYZZZZZ, where XX corresponded to the chromosome (small scaffolds were indicated by 90), Y was 0 or 1 if the gene model was derived from Augustus or StingTie+Transdecoder, respectively, and ZZZZZ followed a consecutive order in the chromosomes.

To validate the new annotation, we used two parallel strategies; search for sequence similarity to a known gene product and search for evidence of expression. We ran BLASTp for all genes in our annotation against three protein sequence pools; UniProt database [62] and the current gene annotation for both *B. floridae* and *B. belcheri.* For all protein sequence pools and each gene independently, we reported a *strong* evidence of sequence similarity for genes with hits fulfilling query length / alignment length > 0.75, subject length / alignment length > 0.75 and e-value < 10^-8^; *weak* evidence for other genes with hits with e-value < 10^-4^; and, *no evidence* for other genes. Gene expression above background noise in *B. lanceolatum* RNA-seq libraries as in [8] was reported as evidence of gene expression for every gene in the new annotation [63]. For every gene, we reported *strong* evidence of gene expression if it was expressed in more than 3 libraries; *weak* evidence if it was expressed in less than 3 libraries and *zero* evidence if expressed in none of the libraries. Of the total 27102 genes, 97.66% (26468) have strong evidence in at least one of the 4 strategies (see complete numbers on annotation validation in Table S5). All genes with reported strong evidence in at least one of the four approaches (3 sequence similarity searches and one gene expression validation) were used for subsequent analysis. This validation method combining sequence similarity to known proteins and gene expression, allows keeping gene models with low similarity, because they evolve fast or are short, while limiting false positives.

### Genome assembly and annotation quality comparison

We compared the quality of BraLan3 with that of other relevant genome references; the previously available assembly for *B. lanceolatum* [8], and the available assemblies for *B. belcheri* [7], *Br. floridae* [9]*, Danio rerio* (GRCz11), *Gallus gallus* (GRCg6a), *Homo sapiens* (GRCh38) and *Mus musculus* (GRCm39). Published *Branchiostoma* genome references were downloaded from NCBI while vertebrate genome references were downloaded from ENSEMBL (release 103). All summary statistics were computed with the final genome assemblies (chromosome level instead of scaffold level if applicable). BUSCO (4.1.4) was run with the metazoan universe (metazoa_odb10) in all cases [64].

### Ortholog and paralog analysis

Genome assemblies, gene annotations and protein sequences for *B. belcheri* [7] and *B. floridae* [9] were downloaded from NCBI genome browser and for *Danio rerio* (GRCz11), *Gallus gallus* (GRCg6a), *Homo sapiens* (GRCh38) and *Mus musculus* (GRCm39) were downloaded from ENSEMBL (release 103). Only BraLan3 genes with strong evidence in at least one validation strategy were used for this analysis (see annotation section). For all species, the sequence of each gene was extracted from genome sequence according to annotated coordinates and was compared to the corresponding protein sequence. The longest gene isoform (transcript) was used in all cases. We filtered out a few genes with non-corresponding annotated sequence-protein sequence pairs mainly a product of annotation errors. We allowed for 10% of mismatches. This filter resulted in a slight reduction of the number of genes used for orthology analysis with respect to the original number of genes in each species gene annotation (see Table S6 for specific numbers). Gene orthology analysis based on protein sequence similarity was performed with Broccoli (version 1.1) [32] using default parameters, DIAMOND (2.0.7.14) [65] and FastTree (2.1.10) [66]. Broccoli groups genes derived from one single gene in the last common ancestor of all considered species in a given orthogroup. Importantly, by using this algorithm, we classified gene duplications preceding chordates’ last common ancestor in different orthogroups and, thus, they were not considered as duplicated in the subsequent analysis. That is, in the orthogroup analysis, only duplications posterior to the last common ancestor of amphioxus and vertebrates were considered. In order to confirm this premise, we used MAFFT (version 7.475, arguments --globalpair --quiet --anysymbol --allowshift) [67] to build multiple sequence alignment of all protein sequences of genes in each orthogroup and RAxML (version v8.2.X, model PROTGAMMAAUTO) [68] to build gene trees for each orthogroup.

We distinguished between single-copy, small-scale duplicated and ohnolog genes as follows. If a given species had more than one gene in a given Broccoli orthogroup, this orthogroup was considered duplicated (non-single-copy) in this species. If at least one amphioxus or vertebrate species presented a duplication in a given orthogroup, this orthogroup was considered as duplicated in the corresponding lineage (amphioxus or vertebrate). Later on, in vertebrates, we differentiated small-scale duplications from ohnologs by classifying all orthogroups containing at least one ohnolog gene derived from the 2R WGDs as ohnolog orthogroups (list of 2R ohnologs from OHNOLOGS version 2.0 for the four vertebrate species [69]). In order to avoid classifying 3R teleost ohnologs (derived from the teleost-specific WGD) as small-scale duplications in vertebrates while focusing in 2R ohnologs, we considered as non-duplicated all orthogroups that, among vertebrates, were only duplicated in zebrafish and that were known to be retained in the 3R WGD (list of 3R ohnologs from OHNOLOGS version 2.0 for zebrafish [69]). Co-occurence of gene duplication status between amphioxus and vertebrate lineages in orthogroups was tested with a hypergeometric test and p-values were corrected with the Bonferroni correction.

### Functional annotation

Human genes belonging to gene ontology (GO) molecular function and biological processes terms were retrieved by unifying iteratively all genes belonging to all child terms of a given GO term [37,38]. This analysis was restricted to human-*B. lanceolatum* orthologous genes, meaning that only *B. lanceolatum* genes with a human ortholog were considered and vice versa*. B. lanceolatum* genes orthologous to a human gene belonging to a given GO term were considered to belong to this same GO term. This same reasoning was applied back to human genes in order to avoid being more restrictive in considering a gene as belonging to a given GO term in human respect to *B. lanceolatum.* That is, if a human gene in a given orthogroup belonged to a given GO term, all the genes in the orthogroup were considered as belonging to this GO term. Only GO terms with a minimum of 50 genes in both humans and *B. lanceolatum* were considered.

### Gene expression

Amphioxus RNA-Seq gene expression data [8] was pseudoaligned to BraLan3 gene annotation with Kallisto [70]using the *--single --rf-stranded -l 180 -s 20 --bias* parameters. Gene abundances were retrieved with tximport [71]. Zebrafish gene expression estimates in several tissues and developmental stages were retrieved from Bgee version 15 [63] using the R package BgeeDB [72]. Transcripts per million (TPM) values were used to express gene expression, allowing between-sample comparison and correcting for transcript length. Boxplots in Figures 4 and S3 are built with R default *boxplot* function parameters without outliers. That is, boxplot’s boxes expand the interquartile range (IQR), while the horizontal line in the box depicts the median. Box plot’s whiskers expand to the last points within 1.5 times the IQR outside the IQR. All points outside that range are considered outliers and not shown. Amphioxus average gene expression in embryonic stages, male gonads, muscle, neural tube, gut, gills and hepatic diverticulum was matched to zebrafish gene expression in embryo, testis, muscle tissue, brain, intestine, pharyngeal gill and liver respectively. Blastula, female gonads and epidermis were excluded from the analysis due to divergent patterns of gene expression in amphioxus. Gene expression specificity in amphioxus adult tissues and embryonic stages was estimated with the Tau index [73].

## Declarations

### Ethics approval and consent to participate

A ripe adult *B. lanceolatum* individual was collected in Argelès-sur-Mer (France) with a specific permission delivered by the Préfet des Pyrénées-Orientales. *B. lanceolatum* is a non-protected species.

### Consent for publication

Not applicable.

### Availability of data and materials

BraLan3 genome reference and annotation and the *B. lanceolatum* whole-genome PacBio sequencing data and chromatin conformation capture data (HiC) we used to build it are available in the European Nucleotide Archive (ENA) under the accession number PRJEB49647. Code used for the analysis available in GitHub (https://github.com/marinabraso/BraLan3) or in Zenodo (doi:10.5281/zenodo.7097385).

### Authors’ contributions

HE provided the original samples. FMar, AE, FMan, MI, IM, RDA, JLGS and JJT performed the BraLan3 assembly and annotation. AE and MRR validated the genome annotation. MBV performed the gene duplication analysis and results. PP and LLT performed the RAG analysis. FMar contributed to the comparative synteny analysis. MBV and MRR wrote the first draft of the manuscript; MBV, DAH, FMar, IM, HE, PP, MI and MRR wrote the final manuscript. MBV produced the figures. MBV, DAH, FMar and MRR contributed to figure visualization design and optimization. All authors contributed to result interpretation and discussion. All authors read and approved the final manuscript.

## Acknowledgements

We thank Kamil S. Jaron for work on an earlier assembly, and discussion of those preliminary results. We acknowledge Professor Robert Waterhouse and all the members of MRR group for valuable discussions. Furthermore, we thank Consolée Aletti and the Genomic Technologies Facility at University of Lausanne for technical contributions. We dedicate this work to the memory of José L. Gómez-Skarmeta.

