## Supplementary figures for "Parallel evolution of amphioxus and vertebrate small-scale gene duplications"

Marina Brasó-Vives et al.

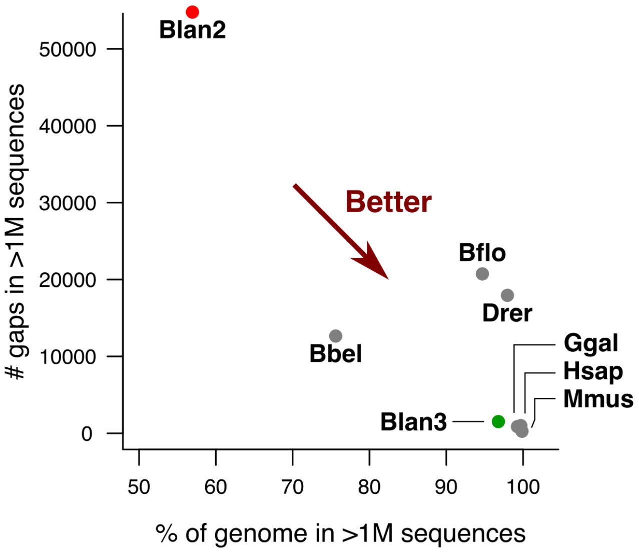


**Figure S1. BraLan3 genome assembly and annotation quality comparison.** Drer, Ggal, Mmus, Hsap, Bflo and Bbel correspond to the latest available genome assemblies for zebrafish, chicken, mouse, human, Florida amphioxus (*B. floridae*) and an Asian amphioxus (*B. belcheri*), respectively (see Methods). Blan2 corresponds to the latest genome reference for the European amphioxus previous to this publication [8] and Blan3 corresponds to BraLan3, the genome reference for the European amphioxus presented in this study. Percentage of the genome assembly sequence in scaffolds (sequences) of length greater than 1 million (M) nucleotides (size of chromosomal magnitude) versus the number of existing sequence gaps in these chromosomal-size sequences.

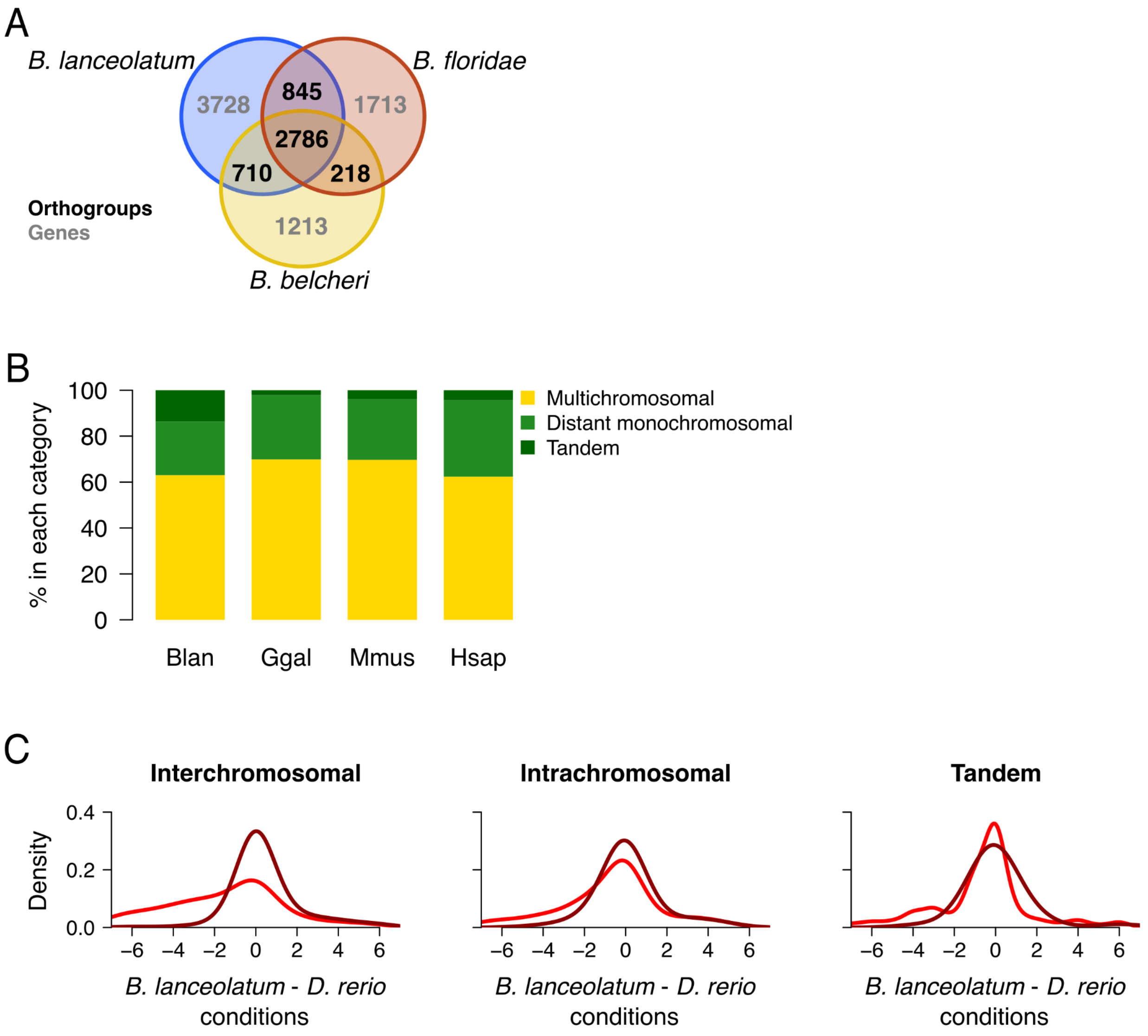


**Figure S2. A.** Venn diagram representing the number of amphioxus specific orthologous groups (black) and amphioxus species-specific genes (gray). **B.** Percentage of small-scale gene duplications that are either multichromosomal, distant monochromosomal and tandem gene duplications in *B. lanceolatum*, chicken, mouse and human (Blan, Ggal, Mmus and Hsap respectively). Zebrafish is excluded from this comparison because of the effect that the 3R WGD in the teleost lineage could have had in the chromosomal localization of small-scale duplications. **C.** Similar to Figure 5 for amphioxus inter-chromosomal, intra-chromosomal distant and tandemly duplicated small-scale duplicates that have single-copy orthologs in zebrafish. For **B** and **C**, Tandem gene duplications are defined as consecutive genes along the chromosomal sequence.

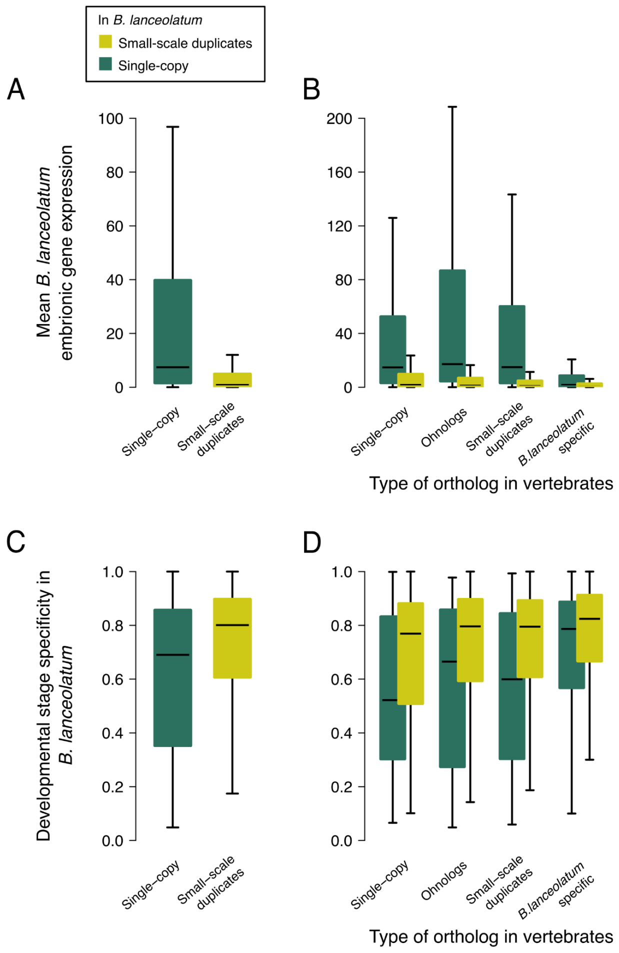


**Figure S3. Embryonic gene expression in single-copy and duplicated genes in *B. lanceolatum*. A.** Box plots showing the distribution of the mean gene expression in transcripts per million (TPM) units across embryonic developmental stages for *B. lanceolatum* single-copy and small-scale duplicated genes. **B.** Same as **A** but dividing *B. lanceolatum* genes according to their vertebrate ortholog (single-copy, ohnolog, small-scale duplicate or *B. lanceolatum* specific). **C-D**. Box plots showing the distribution of developmental stage specificity of *B. lanceolatum* gene expression (Tau statistic) for the same gene groups as **A** and **B**. Tau values range from 0 (ubiquitous expression) to 1 (expression only in 1 tissue).

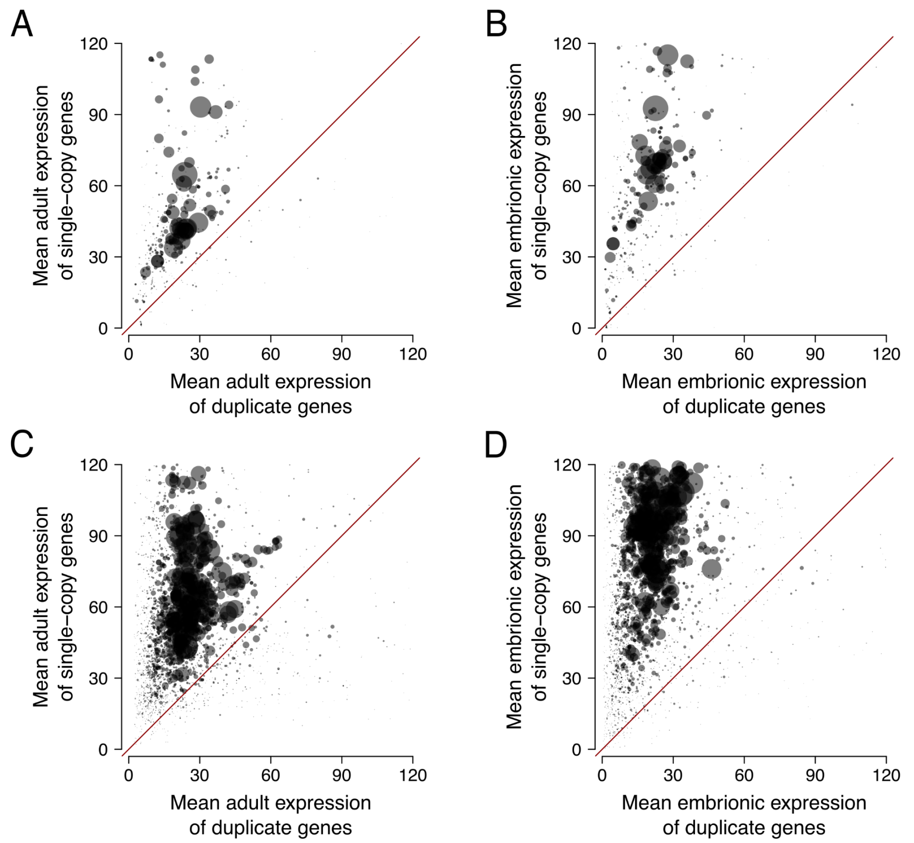


**Figure S4. Lower level of expression of duplicated genes compared to single copy genes within functional categories.** Each point corresponds to a Gene Ontology (GO) term and its size refers to its number of human genes. **A** and **B** show molecular function categories, while **C** and **D** show biological processes categories. **A** and **C** show average expression in adult tissues, while **B** and **D** show average expression in embryonic stages. All points are filled with the same semi-transparent black color, thus opaque regions imply a high point density.

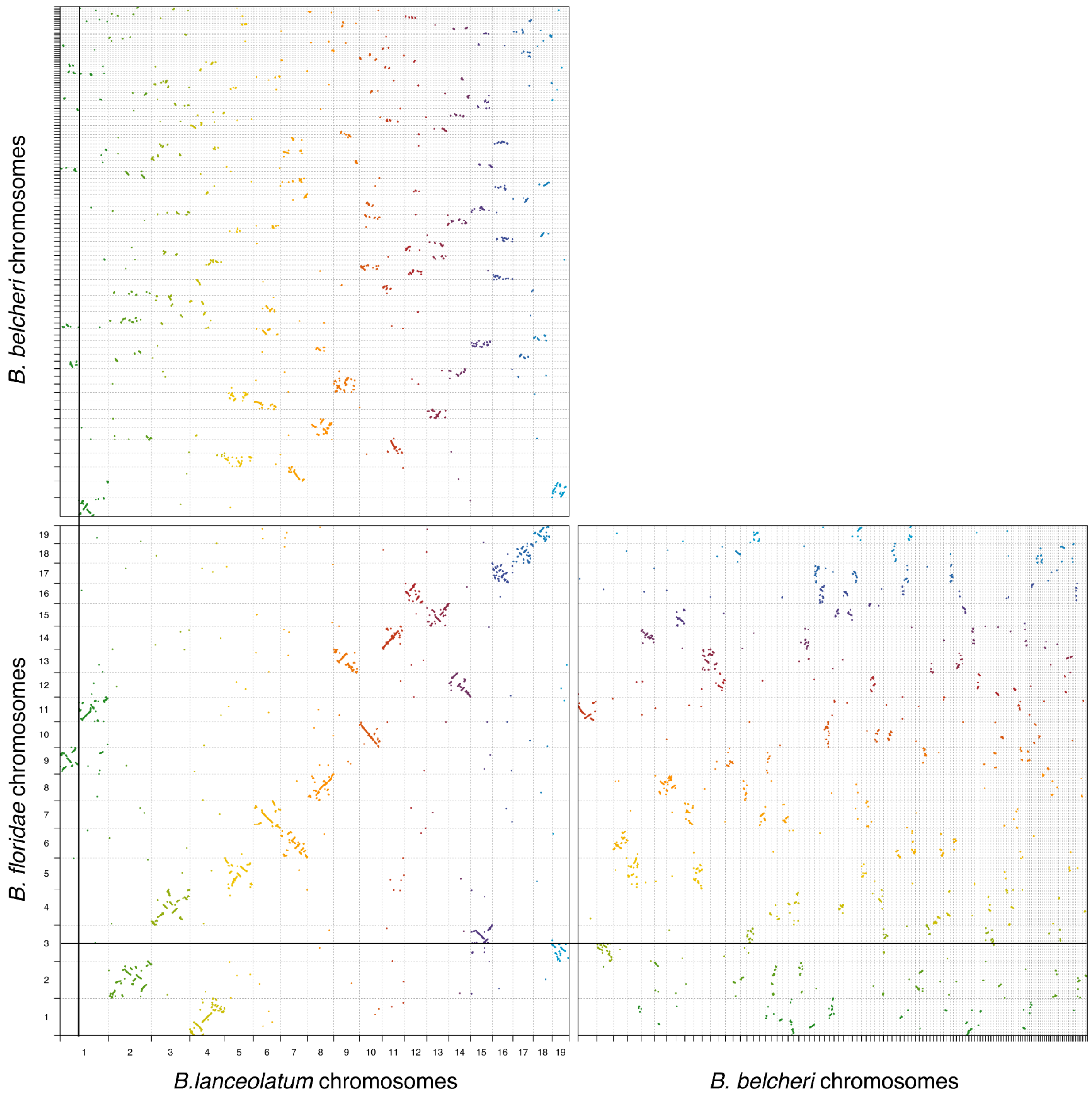


**Figure S5. Gene synteny comparison of *B. lanceolatum* and *B. floridae* with *B. belcheri*.** One-to-one orthologs' synteny conservation between *B. lanceolatum, B. floridae* and *B. belcheri*. Black lines across plots correspond to the fusion/fission points of chromosome 1 in *B. lanceolatum* and chromosome 3 in *B. floridae*. Different colors depict different *B. lanceolatum* chromosomes on the left plots and different *B. floridae* chromosomes on the right plot.

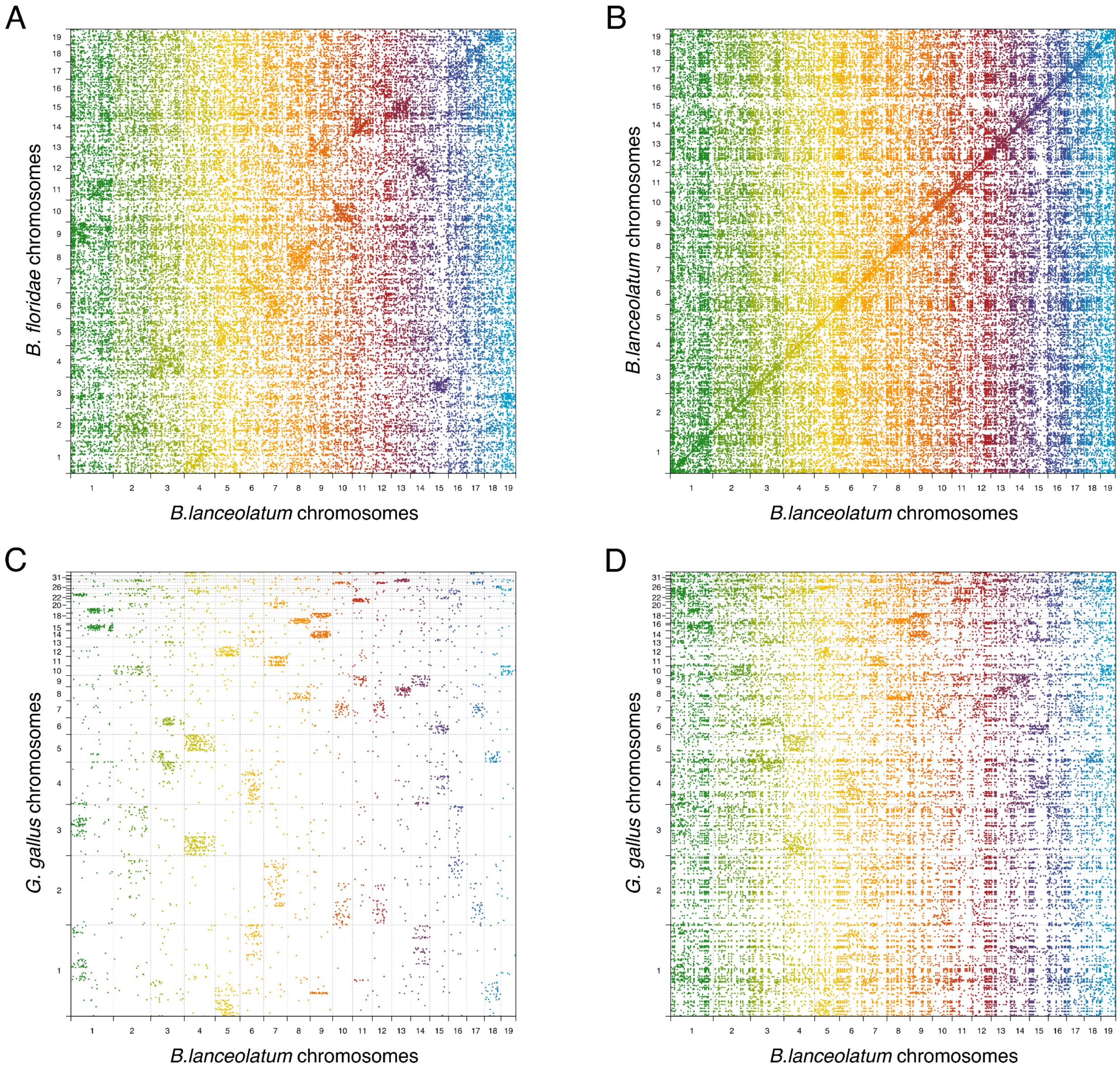


**Figure S6. *B. lanceolatum* gene synteny comparison with *B. floridae* and *G. gallus*. A.** Duplicated genes (many-to-many, many-to-one and one-to-many orthologs) synteny conservation between *B. lanceolatum* and *B. floridae*. **B.** *B. lanceolatum* within species duplicated genes (many-to-many, many-to-one and one-to-many orthologs) synteny conservation. **C.** One-to-one orthologs synteny conservation between *B. lanceolatum* and chicken (*G. gallus*). **D.** Same as in A for *B. lanceolatum* and chicken (*G. gallus*). In all cases (A-C) every gene is represented by its mid-point coordinates in the genome. Different colors depict different *B. lanceolatum* chromosomes.
