## Supplementary note for "Parallel evolution of amphioxus and vertebrate small-scale gene duplications"

Brasó-Vives et al.

### Supplementary note 1: *Proto-RAG*

The recombination-activating genes (RAG1 and RAG2) in jawed vertebrates encode the two subunits of a protein complex essential for V(D)J recombination in immunoglobulin genes. They have their origin in a transposon domestication in vertebrate evolution, a RAGB transposon. An active *Proto-RAG* transposon has been described in the *B. belcheri* genome containing both RAG1 and RAG2 genes flanked by terminal inverted repeats (TIR) (Huang *et al.* 2016). The TIR has a heptamer-spacer-nonamer structure similar to that of the recombination signal sequence (RSS) essential for V(D)J recombination in jawed vertebrates.

We searched for RAGB (or *Proto-RAG*) transposons in *B. lanceolatum* genome, by performing a tBLASTn with the sequences of *Proto-RAG1* (UniProt: A0A185KID9) and *Proto-RAG2* (UniProt: A0A185KIE0) from *B. belcheri* against BraLan3. We encountered three regions containing both *Proto-RAG1* and *Proto-RAG2* blast hits (referred to as sequences 12, 16 and 18 according to the chromosome of their respective location). TIR sequences were found in the flanking sequences of all these regions by performing a BLASTn with known *Proto-RAG* TIR sequences of *B. blecheri* (Tao *et al.* 2020) against these regions' sequences with 10kpb flanking sequences.

To predict completeness or fragmentation of *Proto-RAG* genes in the encountered sequences, we annotated the transposon genes with Softberry FGENESH++ (Solovyev *et al.* 2006). This approach allowed us to see that sequences 12 and 18 seemed complete and potentially active, while sequence 16 appeared incomplete and stopped at the level of *Proto-RAG1*. Sequences 12 and 18 have 98.8% of sequence similarity and present the same structure described for *Proto-RAG* transposon in B. belcheri with which they share 75% of sequence identity in the RAG1 and RAG2 genes sequences. They lack a PHD domain in RAG2 gene sequence, unlike what is seen in jawed vertebrates' RAG2 sequences. The TIR sequences of the three *B. lanceolatum* transposons (including the truncated sequence 16) are identical and show high conservation with other amphioxus *Proto-RAG* TIR sequences (see alignments below).

Finally, we searched BraLan3 for *Miniature Inverted-repeat Transposable Elements* (MITE) sequences containing 5 'and 3' TIRs separated by less than 1000 bp but containing neither RAG 1 and RAG 2. We found 13 MITE sequences in 10 different chromosomes. These results suggest that the *Proto-RAG* transposon is active in the amphioxus genome.


Huang S., X. Tao, S. Yuan, Y. Zhang, P. Li, *et al.*, 2016 Discovery of an Active RAG Transposon Illuminates the Origins of V(D)J Recombination. Cell 166: 102–114. https://doi.org/10.1016/j.cell.2016.05.032

Solovyev V., P. Kosarev, I. Seledsov, and D. Vorobyev, 2006 Automatic annotation of eukaryotic genes, pseudogenes and promoters. Genome Biol. 12.

Tao X., S. Yuan, F. Chen, X. Gao, X. Wang, *et al.*, 2020 Functional requirement of terminal inverted repeats for efficient *ProtoRAG* activity reveals the early evolution of V(D)J recombination. Natl. Sci. Rev. 7: 403–417. https://doi.org/10.1093/nsr/nwz179


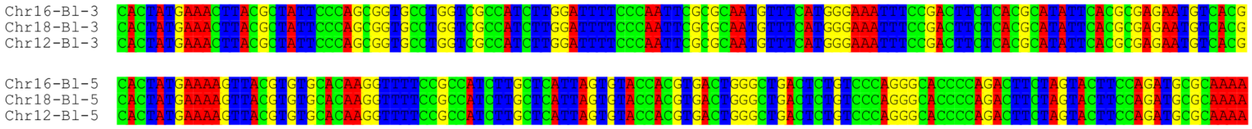


Aligment of the TIR sequences of the tree *Proto-RAG* transposons found in BraLan3 (5' and 3' separately). Both the 3' and the 5' TIR sequences of the tree Proto-RAG tranposons found in *B. lanceolatum* genome are identical in sequence.

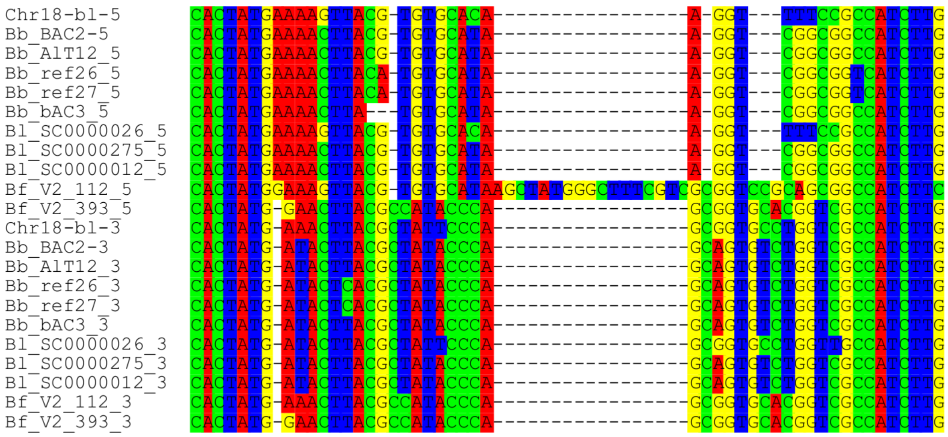


Joined aligment of 5' and 3' TIR sequences of *Proto-RAG* transposons identified in Branchiostoma species. Chr18-bl-5 and Chr18-bl-3 correspond to the 5' and 3' TIR sequences from the ProtoRAG transposon identified here in BraLan3 chromosome 18 (as representatives of the BraLan3 TIR sequences). The other TIR sequences are taken from (Tao *et al.* 2020). Bl, Bb and Bf correspond to B. lanceolatum, B. belcheri and B. floridae respectively while the identification numbers correspond to their insertion scaffolds.
